# Chronic Repeated Predatory Stress Induces Resistance to Quinine Adulteration of Ethanol in Male Mice

**DOI:** 10.1101/677146

**Authors:** Gladys A. Shaw, Maria Alexis M. Bent, Kimaya R. Council, A. Christian Pais, Ananda Amstadter, Jennifer T. Wolstenholme, Michael F. Miles, Gretchen N. Neigh

## Abstract

**Background:** Trauma related psychiatric disorders, such as posttraumatic stress disorder (PTSD), and alcohol use disorder (AUD) are highly comorbid illnesses that separately present an opposing, sex-specific pattern, with increased prevalence of PTSD in females and increased prevalence of AUD diagnoses in males. Likewise, PTSD is a risk factor in the development of AUD, with conflicting data on the impact of sex in the comorbid development of both disorders. Because the likelihood of experiencing more than one traumatic event is high, we aim to utilize chronic repeated predatory stress (CRPS) to query the extent to which sex interacts with CRPS to influence alcohol consumption, or cessation of consumption.

**Methods:** Male (n=16) and female (n=15) C57BL/6J mice underwent CRPS or daily handling for two weeks during adolescence (P35-P49) and two weeks during adulthood (P65-P79). Following the conclusion of two rounds of repeated stress, behavior was assessed in the open field. Mice subsequently underwent a two-bottle choice intermittent ethanol access (IEA) assessment (P90-131) with the options of 20% ethanol or water. After establishing drinking behavior, increasing concentrations of quinine were added to the ethanol to assess the drinking response to adulteration of the alcohol.

**Results:** CRPS increased fecal corticosterone concentrations and anxiety-like behaviors in the open field in both male and female mice as compared to control mice that had not been exposed to CRPS. Consistent with previous reports, we observed a sex difference in alcohol consumption such that females consumed more ethanol per gram of body mass than males. In addition, CRPS reduced alcohol aversion in male mice such that higher concentrations of quinine were necessary to reduce alcohol intake as compared to control mice. CRPS did not alter alcohol-related behaviors in female mice.

**Conclusion:** Collectively, we demonstrate that repeated CRPS can induce anxiety-like behavior in both sexes but selectively influences the response to ethanol adulteration in males.

## 1. Introduction

Traumatic events are highly prevalent, with upwards of 70% of adults worldwide reporting a history of exposure to at least one type of trauma [1] and exposure to serial traumas occuring in as much as 85% of inner city populations [2]. Exposure to trauma increases risk for a number of psychiatric disorders across the internalizing spectrum, including posttraumatic stress disorder (PTSD) [3]. A wealth of data underscore the cumulative effect of trauma, such that higher trauma load across the lifespan is associated with increased risk of psychopathology and severity of symptoms/impairment [4]. Traumatic life events are also associated with increased risk for aberrant drinking and alcohol use disorder (AUD) [5–11], particularly among those with PTSD [12–15]. Given the public health burden of trauma and resulting psychopathology, as well as the clinical importance of comorbid PTSD-AUD (e.g., worse treatment prognosis, increased risk for suicide) [16–20] there exists a need for preclinical models that represent repetitive and complex stress exposures to better understand disease etiology.

The link between mood disorders and ethanol consumption is well studied, with the understanding that prior stress may exacerbate excessive ethanol consumption [21–23]. This connection may be strengthened by the antidepressant effects of ethanol, with its consumption acting as a mechanism of self-medication to mitigate the unwanted psychological effects of stress [24–26]. Early life and adult stress influences the development of alcohol use disorder [27–31], but few studies have considered the potential sex differences in these effects using preclinical models. In regard to mood disorders and alcohol use disorder, males and females exhibit differences in prevalence, with males exhibiting a nearly two fold increased rate of alcohol disorders compared to females [9,32] and females with higher incidences of mood disorders, namely PTSD, such that the prevalence is twice that of males [32–37]. However, studies of comorbidity between PTSD and AUD are inconclusive with regard to sex differences [22]. In order to provide a basis for future mechanistic studies, the current study examined the potential for sex differences to manifest in the phenotypic outcome of a complex repetitive stressor that occurred over two life phases. The goal was not to establish the relative contributions of stress during either individual life phase, but rather to determine the behavioral phenotype following multiple life phase exposures to chronic repeated predatory stress (CRPS) in both sexes.

## 2. Methods

### 2.1 Animals

Juvenile C57BL/6J mice (n = 7-8/group) were purchased from the Jackson Laboratory (Bar Harbor, ME, USA) and arrived at our facilities on postnatal day 22 (**Figure 1A**). Temperature and humidity was maintained at 21°C (± 1°C) and 47%-66% respectively. Mice were pair housed in ventilated rack cages with a sex-matched cage mate in an AAALAC-approved facility. All mice were kept on a 12:12 light:dark cycle (lights on at 0600) with water and Teklad LM-485 7012 standard rodent chow (Envigo, Madison, WI, USA) provided *ad libitum*. On postnatal day 35, mice were randomly assigned into one of two groups (CRPS (male: n= 8, female: n = 7) vs. control (male: n = 8, female: n = 7, **Figure 1B**)). All animal protocols were approved by Virginia Commonwealth University’s Animal Care and Use Committee. All studies were carried out in accordance with the National Institute of Health Guide for the Care and Use of Laboratory Animals.

**Figure 1:**
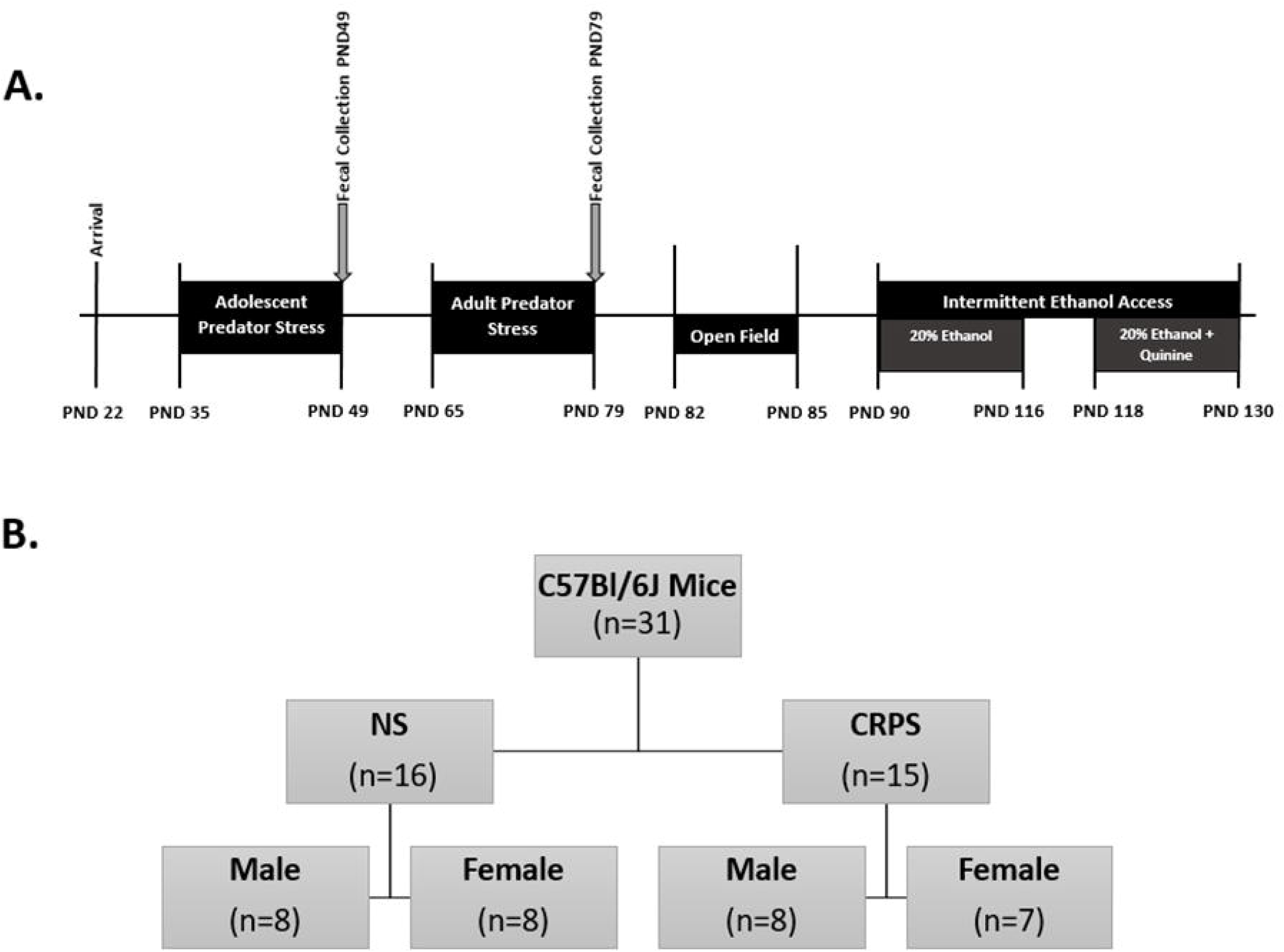
Experimental Design and Timeline. **A)** Experimental timeline. **B)** 31 mice were used in this experiment, 16 males and 15 females. Each sex was separated equally into non stress and chronic repeated predatory stress (CRPS) groups. Mice subject to CRPS were chosen at random at PND 35 before the beginning of CRPS.

### 2.2 Chronic Repeated Predatory Stress (CRPS)

As previously described [38–40], predatory stress was completed daily for a total of fifteen days in adolescence (post-natal day [PND] 35-49) and fifteen days in adulthood (PND 65-79) (**Figure 1A**). Stress during adolescence is well-documented to cause long-lasting changes in physiology and behavior [38,40–50] and repeated exposures to trauma are pervasive in populations at high risk for trauma-related pathophysiology [2]. Consistent with previous publications [38–40], predatory stress was comprised of a thirty-minute exposure of the mice to a Long Evans rat. All Long Evans rats were retired male breeders, restricted to 15g rodent chow per day, and had reduced cage changes from twice to once a week throughout the duration of the stress to increase aggression. During their light cycle, mice were placed in a dwarf hamster ball measuring five inches in diameter and then placed in the home cage of the rat (Lee’s Aquarium & Pet Products, San Marcos, CA, USA, Cat. #20198 and #20193).

### 2.3 Fecal corticosterone assessment

Fecal boli were collected during the study (**Figure 1A**) in order to assess alterations in corticosterone concentrations. Collections were obtained on PND 49, following either the 15^th^ exposure to stress or 15^th^ exposure to handling, and PND 79, following the 30^th^ exposure to stress or the 30^th^ exposure to handling. All collections occurred during the light cycle a minimum of two hours prior to the onset of the dark cycle. Mice were briefly placed in an empty mouse cage that had been cleaned with 70% ethanol. Mice remained in the cage for approximately two minutes per collection. Two fecal boli per mouse were collected using a clean, irradiated wooden toothpick, placed in a clean 2.0mL micro centrifuge tube. Freshly collected fecal boli were stored on dry ice for transport and moved to a −80°C freezer for long term storage. Fecal samples were processed according to methods described in Bardi et al., 2011 [51]. Samples were thawed for one hour at room temperature preceding corticosterone assessment. Thawed fecal samples were weighed and collected in glass test tubes with 500μL of 100% methanol per 45mg fecal coli. Methanol volume was scaled proportionally to the weight of fecal matter collected. Glass tubes were covered in Parafilm, vortexed for approximately 30s to homogenize the sample, and centrifuged at 2500xg for 10 minutes at 23°C. The supernatant was removed and diluted 1:20 in assay buffer included in the Enzo Corticosterone ELISA kit (Cat. No. ADI-901-097). Assay was completed in duplicate according to manufacturer’s instructions. Plate was read at both 405nm and 535nm using an automated plate reader to determine concentration of metabolized corticosterone at each time point.

### 2.4 Estrus Assessment

On postnatal day 79, estrus was measured in all female mice using a visual assessment based on the characterization found in Byers et al., 2012 [52]. Estrus tracking continued daily throughout behavioral assessments. Males were handled in a similar fashion to mitigate the confound of differential handling between the sexes.

### 2.5 Open Field Test

The open field test was used to evaluate both locomotor activity and anxiety-like behavior [53–55]. Testing was conducted at least three days after the final stressor using methods similar to those previously described [38]. Equal numbers of mice from each group were assessed on each of the testing days. Each mouse was only assessed one time. Three to four hours into their light cycle, mice were placed in a 13.5” x 13.5” white bottom polyethylene square box with 16” high white walls and permitted to explore for ten minutes. Activity was recorded using an overhead camera, and metrics were assessed using EthoVision XT13 Software (Noldus Technologies, Leesburg, VA, USA). Percentage of time spent in the center of the arena, velocity, percent of total time spent moving, and total distance traveled were assessed.

### 2.6 Intermittent Ethanol Access (IEA)

Eight days after the last CRPS exposure, mice were transferred from ventilated rack cages to static home cages in preparation for ethanol assessments. Mice were given three days (PND 87-89) to habituate to their new cage. All mice were individually housed to ensure accuracy in assessment of drinking volumes. Two bottle choice intermittent ethanol exposure was executed as described [56]. Over the course of four weeks, mice were given a choice between 20% (v/v) ethanol made from 100% ethanol and tap water, or unaltered tap water in sipper tubes, constructed from 10mL serological pipettes and fitted with metal ball-bearing sipper tubes. All mice were given access to water or 20% ethanol three days a week, every Mon-Wed-Fri, *ad libitum*, on a twenty-four hour on/off cycle, one to two hours before their dark cycle [57]. Starting and ending volumes were noted and used to determine consumption during the twenty-four-hour time period. Sipper tube volumes, not weights, were used to assess fluid consumption and to remove the confound of fluid loss due to the removal of the tubes from the cage. Two empty control cages were used to assess fluid loss due to leaking and evaporation of both water and 20% ethanol. Starting and ending volumes of the tubes in the control cages were noted during each twenty-four hour time period, averaged, and subtracted from the consumption values in each mouse cage.

### 2.8 Quinine Exposure (QuiA)

To determine if aversion-resistant alcohol drinking was present, quinine, a bitter tastant, was incorporated into the 20% ethanol as previously described [57]. Quinine exposure is commonly used to determine aversion-resistant drinking behavior, as it produces a negative effect linked to ethanol consumption, a defining factor of the human definition of AUD [57]. Increasing concentrations of quinine were added into 20% ethanol, at the concentrations of 5mg/L, 10mg/L, 50mg/L, 100mg/L, 150mg/L, and 200mg/L over the course of twelve days. Mice were given twenty-four hours to consume either water or 20% ethanol with the specified concentration of quinine in the same two-bottle choice model described above. Methods to assess ethanol-quinine consumption and preference were measured identically to the initial IEA measurements.

### 2.9 Statistical analysis

GraphPad Prism 8.0.2 for Windows (GraphPad Software, La Jolla, CA) was used for assessment of all subsequent analyses. An alpha value of 0.05 was used in all cases. Fecal corticosterone data were analyzed using a repeated measures three-way ANOVA with the factors of time, sex, and stress. Behavioral data were analyzed using a two-way ANOVA with the factors of sex and stress. Significant interactions were further assessed using Tukey’s post hoc analysis. Consistent with a significant sex difference in ethanol consumption reported throughout the literature for C57BL/6J mice [58–60], IEA data were analyzed first using a three-way repeated measures ANOVA with the factors time, sex, and stress. Follow up analyses were separated between male and female and run using a two-way repeated measures ANOVA with the factors time and stress to determine 20% ethanol intake by volume, ethanol preference, or total fluid volume consumed. When interactions were present, post hoc analyses were completed using Tukey’s post hoc analysis.

## 3. Results

### 3.1 Fecal corticosterone

Exposure to CRPS increased corticosterone concentrations, as compared to handled controls (F_(1, 27)_ = 26.37, p < 0.0001; **Figure 2**). In addition, corticosterone concentrations were higher in female than male mice (F_(1, 27)_ = 13.15, p = 0.0012). There was no effect of time on corticosterone concentrations such that values were similar within each group between PND 49 and PND 79 (p > 0.05).

**Figure 2:**
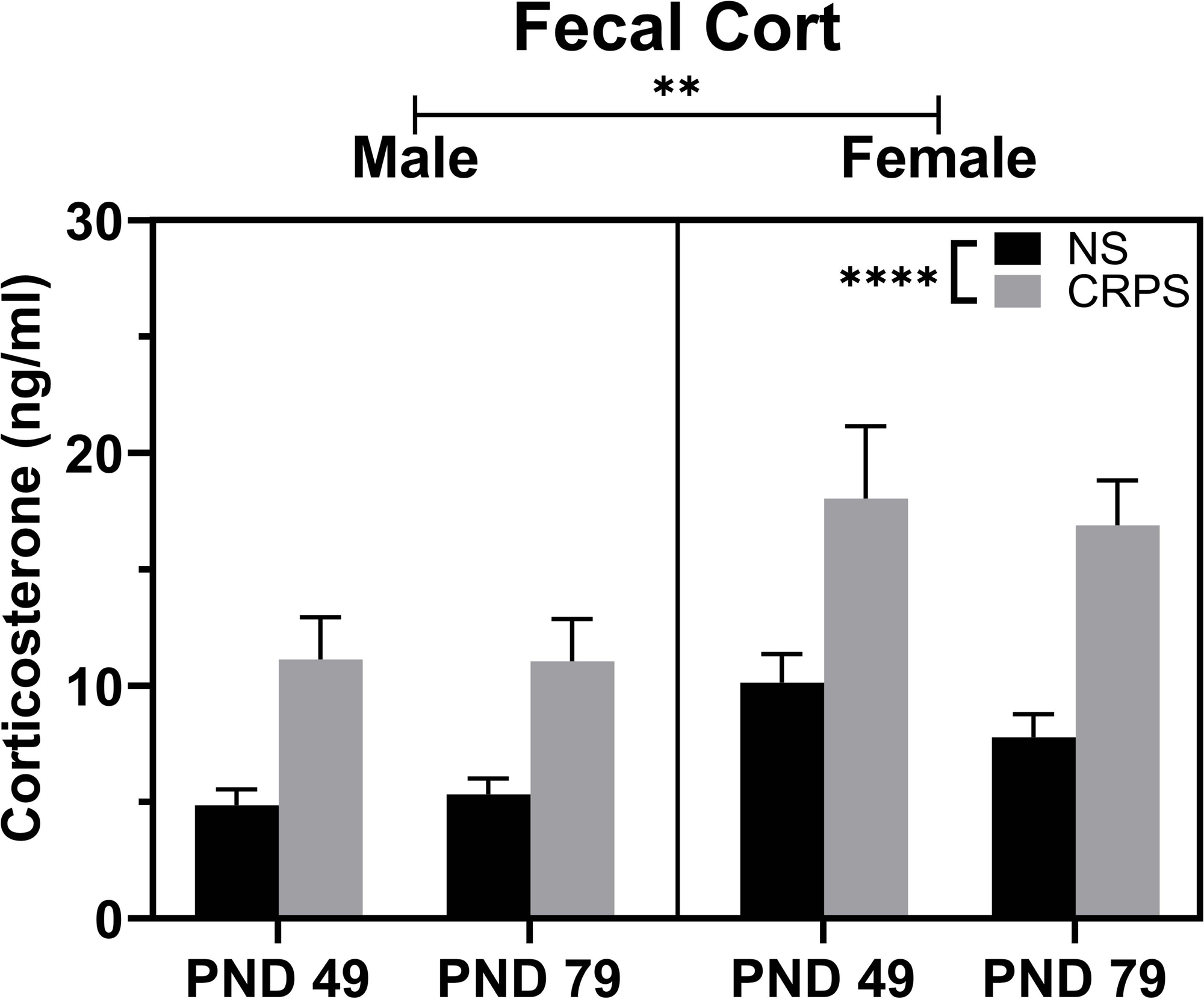
CRPS Increases Corticosterone Concentrations. Corticosterone concentrations collected following the 15^th^ (PND 49) and 30^th^ (PND 79) exposure to handling (control) or chronic repeated predatory stress (CRPS group) were collected from fecal boli. Corticosterone concentrations were higher in the CRPS groups of both sexes at both timepoints and females had higher corticosterone concentrations in fecal boli than males. Reported values depict mean ± SEM. #p = 0.06, **p < 0.01, ***p < 0.0001.

### 3.2 Open Field

Exposure to CRPS reduced overall time in the center of the arena (F_(1,27)_ = 8.780; p = 0.0063, **Figure 3A**) when compared to the non-stressed controls. There was no effect of sex on time spent in the center of the arena (p > 0.05). Mice with a history of CRPS displayed signs of hyperactivity, as these mice spent more time moving (F_(1,27)_ = 18.47, p = 0.0002), covered a greater distance (F_(1,27)_ = 15.22, p = 0.0006), and moved at a higher speed (F_(1,27)_ = 14.54, p = 0.0007) than non-stressed control mice (**Figure 3B-D**). Three of the seven female mice with a history of CRPS were in estrus during the time of the open field assessment. Pearson’s correlation analysis showed no significant effect of estrus stage in open-field performance within this group (p > 0.05). No non-stressed control females were in estrus at the time of testing.

**Figure 3:**
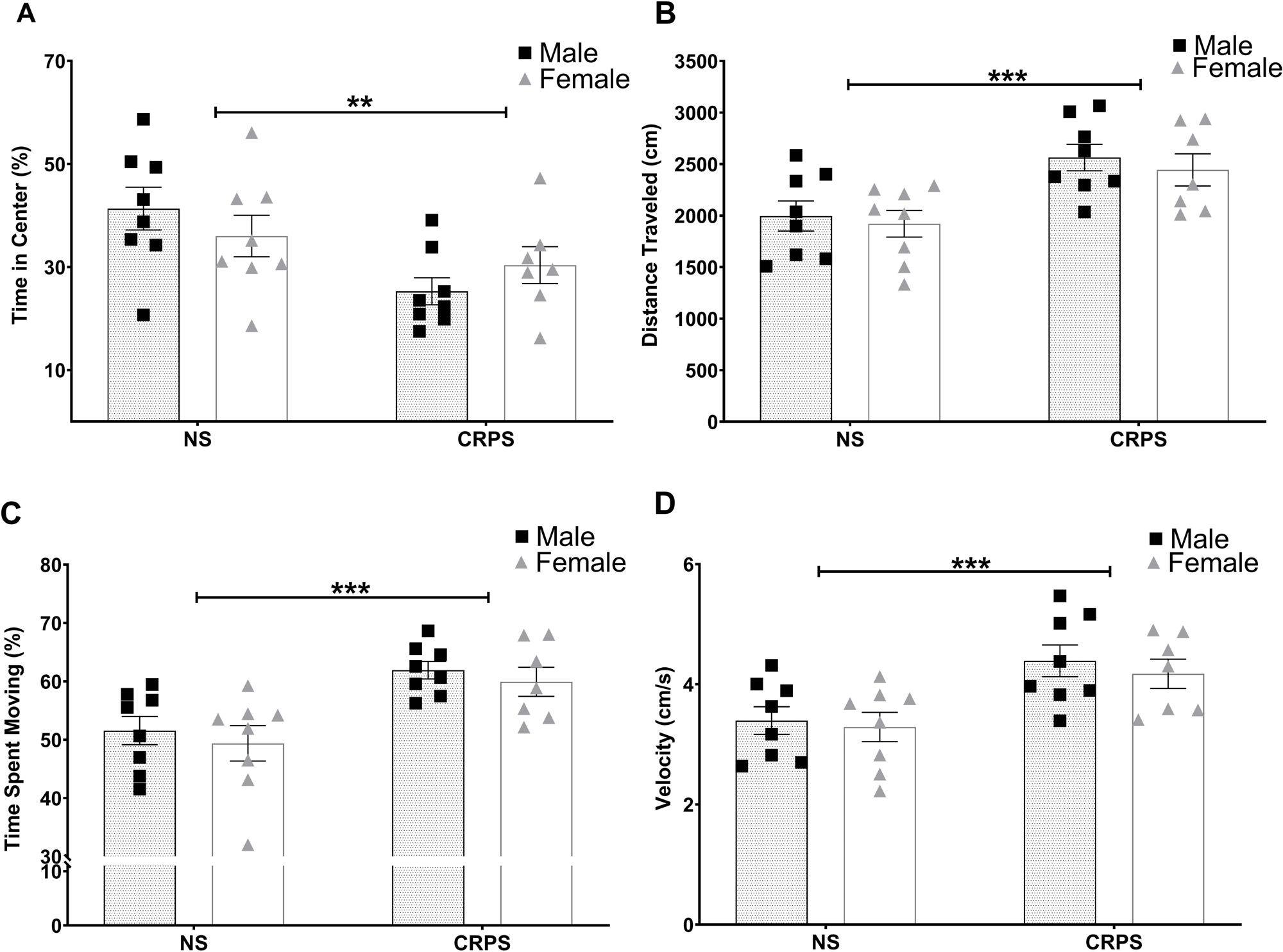
CRPS increases anxiety-like behaviors in the open field test. **A)** Chronic repeated predatory stress (CRPS) decreased time spent in the center regardless of sex when compared to controls. **B-D)** Likewise, a history of CRPS increased locomotor activity in the open field. Reported values depict mean ± SEM. *p < 0.05, **p < 0.01, ***p < 0.001.

### 3.4 Intermittent Ethanol Access

As expected with the IEA model and consistent with the literature, the mice significantly increased their ethanol consumption (F_(5.711, 154.2)_ = 21.51, p < 0.0001; **Figure 4A**) and preference (F_(4.691, 126.7)_ = 13.77, p < 0.0001; **Figure 4B**) over time (Hwa et al., 2011). Female mice consumed more alcohol (adjusted for body mass) than male mice (F_(1,27)_ = 43.87, p < 0.0001), as expected based on previous work [60]. A history of CRPS did not significantly increase either alcohol consumption (p > 0.05) or preference for the 20% alcohol solution (p > 0.05). Although total fluid intake differed by sex (F_(1,27)_= 37.71, p < 0.0001) and time (F_(11, 297)_ = 19.41, p < 0.0001), consistent with the sex difference in consumption reported above, previous exposure to CRPS did not alter total fluid intake (p > 0.05), suggesting differences in ethanol intake and preference are not due to differences in total fluid intake following CRPS exposure (data not shown).

**Figure 4:**
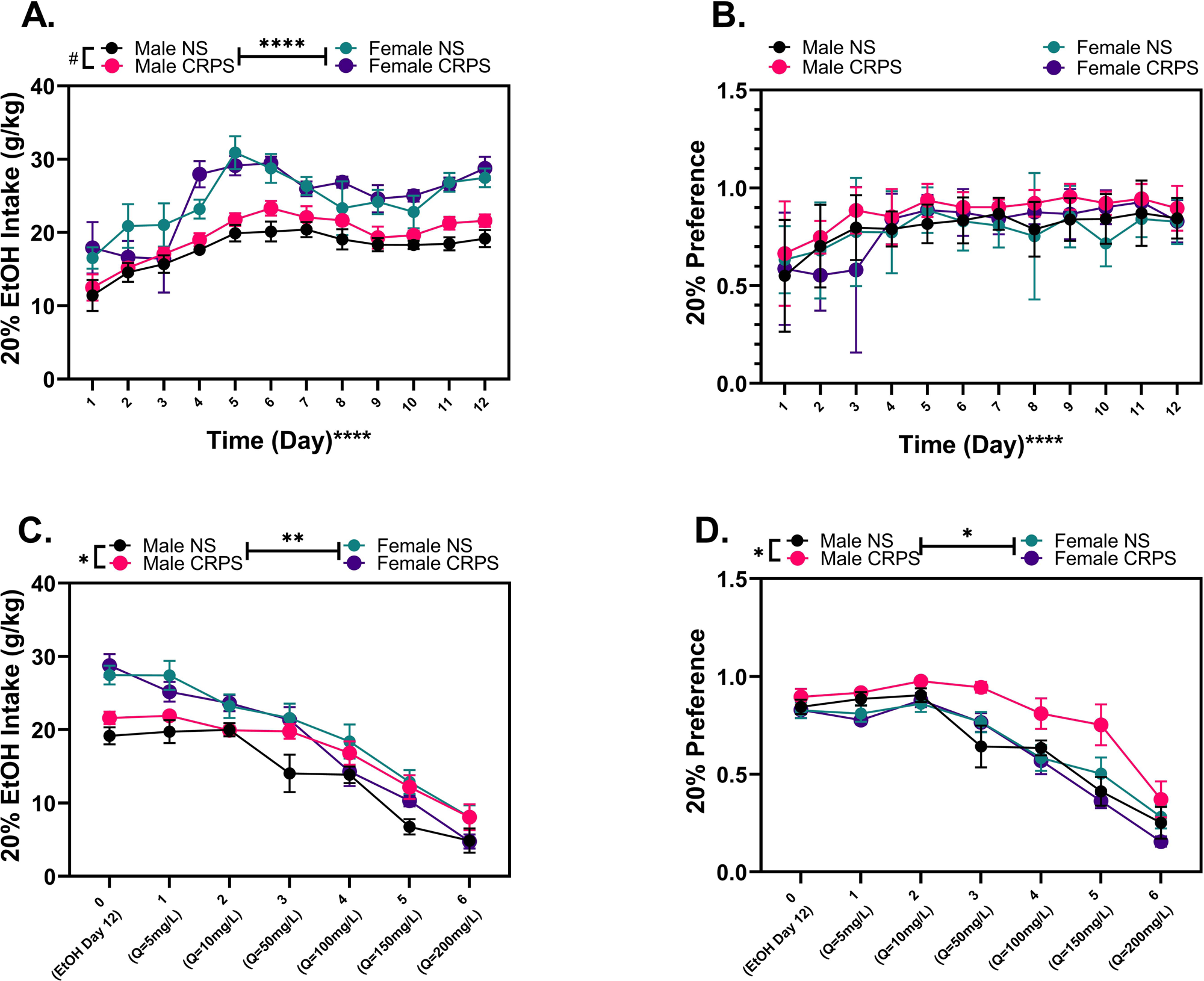
CRPS does not alter alcohol intake. **A)** Baseline ethanol intake by weight shows a sex difference between ethanol consumption (p < 0.0001), with females consuming more ethanol by weight than males regardless of chronic repeated predatory stress (CRPS) history. **B)** There were no baseline difference in ethanol preference in terms of sex or CRPS history. **C)** Males with a history of CRPS consumed more quinine adulterated ethanol than male controls. Within females, there was no significant difference in quinine adulterated ethanol consumption between the CRPS backgrounds. **D)** Males with a history of CRPS had a higher quinine adulterated 20% ethanol preference than control males. Control males significantly decreased quinine adulterated ethanol consumption at 100mg/L quinine; whereas, males with a history of CRPS significantly decreased at 200mg/L quinine adulterated ethanol. Within females, there was no significant difference in quinine adulterated ethanol preference between the controls and mice with a history of CRPS. Reported values depict mean ± SEM. *p < 0.05, **p < 0.01, ***p < 0.001, ****p < 0.0001.

### 3.5 Quinine

Exposure to CRPS delayed the impact of quinine adulteration (QuiA) on alcohol intake such that higher concentrations of quinine were necessary to reduce drinking behaviors (main effect of time: F_(4.368,117.9)_ = 126.6, p < 0.0001). Sex (F_(1,27)_ = 11.62, p = 0.0021), but not CRPS (p > 0.05), influenced intake following quinine adulteration (**Figure 4C**). The impact of sex on intake after quinine adulteration was specific to the male groups (F_(1,14)_ = 6.448, p = 0.0236). Males with a history of CRPS consumed more quinine adulterated ethanol than non-stressed males. Preference for quinine-adulterated ethanol (**Figure 4D**) was impacted by sex (F_(1,27)_ = 5.792, p = 0.0232), an interaction between sex and CRPS (F_(1,27)_ = 6.889, p = 0.0141) and an interaction between quinine concentration, sex, and CRPS (F_(6,162)_ = 2.752, p = 0.0142). Follow up analyses in males demonstrated that there was a main effect of CRPS (F_(1,14)_ = 8.406, p = 0.0117) and a significant interaction between quinine dose and CRPS (F_(6,84)_ = 2.438, p = 0.0320) on preference. Non-stressed males consumed significantly less quinine laced ethanol when quinine levels reached 100mg/L (p = 0.0133) compared to baseline; whereas, males with a history of CRPS did not significantly decrease consumption until quinine levels reached 200mg/L (p = 0.0049). There was no significant difference in ethanol consumption or preference between females regardless of CRPS background (p > 0.05) or non-stressed control male and female mice (p > 0.05).

## 4. Discussion

CRPS exposure precipitates resistance to quinine adulteration of ethanol in male, but not female, mice. Although drinking behavior was only impacted in males with a history of CRPS, both males and females exhibited behavioral and neuroendocrine alterations in response to the repeated predatory stressor exposures demonstrating that both sexes were impacted by the stressors and suggesting that males may be selectively sensitive to stress effects on ethanol-related behaviors.

One of the hallmarks of AUD is the continuance of drinking despite pervasive negative consequences [57,62]. In order to assess conflict-resistant alcohol consumption we used the previously established method of adulteration of the ethanol solution with quinine, a bitter tastant [63]. Despite similar impacts on anxiety-like behavior, the effects of CRPS on ethanol consumption were disproportionately represented in male mice. Although control males, and females regardless of stress history, reduced QuiA consumption, similar to a previous report of no impact of sex on QuiA consumption [64], male mice that had a history of CRPS demonstrated conflict-resistant alcohol drinking. Most remarkably, a history of CRPS caused male mice to be resistant to QuiA such that they continued preferentially choosing to consume the 20% ethanol solution over water until 2x the quinine was added to the solution that was necessary to dissuade drinking in male controls. A similar effect has been demonstrated previously through the use of repeated cycles of alcohol exposure [65] but has not been reported following a chronic stress paradigm in mice. However, similar effects of exposure to a predator stimulus have been reported to increase compulsive-like drinking of QuiA in a subset of male rats, suggesting that the impact of predatory stress may generalize across species [21]. This disassociation in anxiety-like behavior and ethanol consumption behaviors may provide a model system in which to assess the neural underpinnings of CRPS exposure on alcohol behaviors with high translational potential given the role of drinking despite consequences and impairment as a cardinal symptom of AUD.

The utilization of this CRPS model to induce stress not only produces robust, long lasting effects in mice, but also, due to the nature of the stressor, is less likely to give rise to sex-specific differences in affective-like behaviors, unlike other models of predation stress, such as predator odor [66,67]. The data presented in this manuscript extend previous findings [38–40] to demonstrate that predation stress beginning in adolescence alters anxiety-like behaviors in both male and female mice. Importantly, the behavioral assessments reported here were conducted prior to ethanol consumption and do not reflect an impact of ethanol intoxication or withdrawal [68]. The model of CRPS provides a system in which divergences in manifestation of the consequences of a history of chronic stress exposure can be evaluated as both sexes demonstrate a measurable physiological and behavioral change in response to the exposures.

In summary, these data demonstrate that CRPS increases corticosterone concentrations in male and female mice. Moreover, a history of CRPS increases behaviors consistent with an anxiety-like phenotype in both males and females. Despite the similarities in behaviors and neuroendocrine function in both sexes, only male mice with a history of CRPS exhibited resistance to quinine adulteration, which was not observed in females. Mechanistic insight into sex differences in alcohol behaviors may be gained by analysis of the neural underpinnings of the phenotypic differences reported here.

## Acknowledgements

This research was funded by the Specialized Alcohol Research Centers grant (P50) through the National Institutes of Alcohol Abuse and Alcoholism (NIAAA), awarded to Michael F. Miles, MD/PhD (AA022537) and the National Institutes of Health NIAAA R01 grant awarded to Jennifer T. Wolstenholme, PhD (AA026347).

## References

[1] C. Benjet, E. Bromet, E.G. Karam, R.C. Kessler, K.A. McLaughlin, A.M. Ruscio, V. Shahly, D.J. Stein, M. Petukhova, E. Hill, J. Alonso, L. Atwoli, B. Bunting, R. Bruffaerts, J.M. Caldas-de-Almeida, G. de Girolamo, S. Florescu, O. Gureje, Y. Huang, J.P. Lepine, N. Kawakami, V. Kovess-Masfety, M.E. Medina-Mora, F. Navarro-Mateu, M. Piazza, J. Posada-Villa, K.M. Scott, A. Shalev, T. Slade, M. ten Have, Y. Torres, M.C. Viana, Z. Zarkov, K.C. Koenen, The epidemiology of traumatic event exposure worldwide: results from the World Mental Health Survey Consortium, Psychol. Med. 46 (2016) 327–343. doi:10.1017/S0033291715001981.

[2] C.F. Gillespie, B. Bradley, K. Mercer, A.K. Smith, K. Conneely, M. Gapen, T. Weiss, A.C. Schwartz, J.F. Cubells, K.J. Ressler, Trauma Exposure and Stress-Related Disorders in Inner City Primary Care Patients, Gen Hosp Psychiatry. 31 (2009) 505–514. doi:10.1016/j.genhosppsych.2009.05.003.

[3] R.C. Brown, E.C. Berenz, S.H. Aggen, C.O. Gardner, G. Knudsen, R.-K. Ted, K.S. Kendler, A.B. Amstadter, Trauma exposure and Axis I psychopathology: A cotwin control analysis in Norwegian young adults., Psychol. Trauma Theory Res Pr. Policy. 6 (2014) 652. doi:10.1037/a0034326.

[4] E.G. Karam, M.J. Friedman, E.D. Hill, R.C. Kessler, K.A. McLaughlin, M. Petukhova, L. Sampson, V. Shahly, M.C. Angermeyer, E.J. Bromet, G. De Girolamo, R. De Graaf, K. Demyttenaere, F. Ferry, S.E. Florescu, J.M. Haro, Y. He, A.N. Karam, N. Kawakami, V. Kovess-Masfety, M.E. Medina-Mora, M.A.O. Browne, J.A. Posada-Villa, A.Y. Shalev, D.J. Stein, M.C. Viana, Z. Zarkov, K.C. Koenen, Cumulative traumas and risk thresholds: 12-month ptsd in the world mental health (WMH) surveys, Depress. Anxiety. 31 (2014) 130–142. doi:10.1002/da.22169.

[5] C. Erbes, J. Westermeyer, B. Engdahl, E. Johnsen, {Post-Traumatic} Stress Disorder and Service Utilization in a Sample of Service Members from Iraq and Afghanistan, Mil Med. 172 (2007) 359–363. doi:10.7205/milmed.172.4.359.

[6] C.W. Hoge, C.A. Castro, S.C. Messer, D. McGurk, D.I. Cotting, R.L. Koffman, Combat duty in Iraq and Afghanistan, mental health problems and barriers to care., US. Army Med. Dep. J. 351 (2004) 7–17. doi:10.1056/nejmoa040603.

[7] C.W. Hoge, J.L. Auchterlonie, C.S. Milliken, Mental health problems, use of mental health services, and attrition from military service after returning from deployment to Iraq or Afghanistan, J. Am. Med. Assoc. 295 (2006) 1023–1032. doi:10.1001/jama.295.9.1023.

[8] I.G. Jacobson, M.A.K. Ryan, T.I. Hooper, T.C. Smith, P.J. Amoroso, E.J. Boyko, G.D. Gackstetter, T.S. Wells, N.S. Bell, Alcohol use and alcohol-related problems before and after military combat deployment, JAMA - J. Am. Med. Assoc. 300 (2008) 663–675. doi:10.1001/jama.300.6.663.

[9] D.G. Kilpatrick, K.J. Ruggiero, R. Acierno, B.E. Saunders, H.S. Resnick, C.L. Best, Violence and risk of PTSD, major depression, substance abuse/dependence, and comorbidity: Results from the national survey of adolescents, J. Consult. Clin. Psychol. 71 (2003) 692–700. doi:10.1037/0022-006X.71.4.692.

[10] T.C. Smith, M.A.K. Ryan, D.L. Wingard, D.J. Slymen, J.F. Sallis, D. Kritz-Silverstein, New onset and persistent symptoms of post-traumatic stress disorder self reported after deployment and combat exposures: Prospective population based US military cohort study, Bmj. 336 (2008) 366–371. doi:10.1136/bmj.39430.638241.AE.

[11] K.M. Wright, A.H. Huffman, A.B. Adler, C.A. Castro, Psychological Screening Program Overview, Mil Med. 167 (2002) 853–861. doi:10.1093/milmed/167.10.853.

[12] M. Head, L. Goodwin, F. Debell, N. Greenberg, S. Wessely, N.T. Fear, Post-traumatic stress disorder and alcohol misuse: comorbidity in UK military personnel, Soc. Psychiatry Psychiatr. Epidemiol. 51 (2016) 1171–1180. doi:10.1007/s00127-016-1177-8.

[13] R.C. Kessler, A. Sonnega, E. Bromet, M. Hughes, C.B. Nelson, Posttraumatic Stress Disorder in the National Comorbidity Survey, Arch. Gen. Psychiatry. 52 (1995) 1048–1060. doi:10.1001/archpsyc.1995.03950240066012.

[14] I.L. Petrakis, R. Rosenheck, R. Desai, Substance use comorbidity among veterans with posttraumatic stress disorder and other psychiatric illness, Am. J. Addict. 20 (2011) 185–189. doi:10.1111/j.1521-0391.2011.00126.x.

[15] L. Sampson, G.H. Cohen, J.R. Calabrese, D.S. Fink, M. Tamburrino, I. Liberzon, P. Chan, S. Galea, Mental Health Over Time in a Military Sample: The Impact of Alcohol Use Disorder on Trajectories of Psychopathology After Deployment, J. Trauma. Stress. 28 (2015) 547–555. doi:10.1002/jts.22055.

[16] C. Blanco, Y. Xu, K. Brady, G. Pérez-Fuentes, M. Okuda, S. Wang, Comorbidity of posttraumatic stress disorder with alcohol dependence among US adults: Results from national epidemiological survey on alcohol and related conditions, Drug Alcohol Depend. 132 (2013) 630–638. doi:10.1016/j.drugalcdep.2013.04.016.

[17] J.C. Ipser, D. Wilson, T.O. Akindipe, C. Sager, D.J. Stein, Pharmacotherapy for anxiety and comorbid alcohol use disorders, Cochrane Database Syst. Rev. 2017 (2015) CD007505. doi:10.1002/14651858.CD007505.pub2.

[18] J.P. Read, P.J. Brown, C.W. Kahler, Substance use and posttraumatic stress disorders: Symptom interplay and effects on outcome, Addict. Behav. 29 (2004) 1665–1672. doi:10.1016/j.addbeh.2004.02.061.

[19] S.M. Rojas, S. Bujarski, K.A. Babson, C.E. Dutton, M.T. Feldner, Understanding PTSD comorbidity and suicidal behavior: Associations among histories of alcohol dependence, major depressive disorder, and suicidal ideation and attempts, J. Anxiety Disord. 28 (2014) 318–325. doi:10.1016/j.janxdis.2014.02.004.

[20] D. Shorter, J. Hsieh, T.R. Kosten, Pharmacologic management of comorbid post-traumatic stress disorder and addictions, Am. J. Addict. 24 (2015) 705–712. doi:10.1111/ajad.12306.

[21] S. Edwards, B.B. Baynes, C.Y. Carmichael, E.R. Zamora-Martinez, M. Barrus, G.F. Koob, N.W. Gilpin, Traumatic stress reactivity promotes excessive alcohol drinking and alters the balance of prefrontal cortex-amygdala activity., Transl. Psychiatry. 3 (2013) e296. doi:10.1038/tp.2013.70.

[22] N.W. Gilpin, J.L. Weiner, Neurobiology of comorbid post-traumatic stress disorder and alcohol-use disorder, Genes, Brain Behav. 16 (2017) 15–43. doi:10.1111/gbb.12349.

[23] G.F. Koob, M. Le Moal, Drug addiction, dysregulation of reward, and allostasis, Neuropsychopharmacology. 24 (2001) 97–129. doi:10.1016/S0893-133X(00)00195-0.

[24] A.H. Putman, A.R. Wolen, J.L. Harenza, R.K. Yordanova, B.T. Webb, E.J. Chesler, M.F. Miles, Identification of quantitative trait loci and candidate genes for an anxiolytic-like response to ethanol in BXD recombinant inbred strains, Genes, Brain Behav. 15 (2016) 367–381. doi:10.1111/gbb.12289.

[25] S.A. Wolfe, E.R. Workman, C.F. Heaney, F. Niere, S. Namjoshi, L.P. Cacheaux, S.P. Farris, M.R. Drew, B. V. Zemelman, R.A. Harris, K.F. Raab-Graham, FMRP regulates an ethanol-dependent shift in GABA B R function and expression with rapid antidepressant properties, Nat. Commun. 7 (2016) 12867. doi:10.1038/ncomms12867.

[26] S.A. Wolfe, S.P. Farris, J.E. Mayfield, C.F. Heaney, E.K. Erickson, R.A. Harris, R.D. Mayfield, K.F. Raab-Graham, Ethanol and a rapid-acting antidepressant produce overlapping changes in exon expression in the synaptic transcriptome, Neuropharmacology. 146 (2019) 289–299. doi:10.1016/j.neuropharm.2018.11.007.

[27] C. Evren, V. Sar, E. Dalbudak, R. Cetin, M. Durkaya, B. Evren, S. Celik, Lifetime PTSD and quality of life among alcohol-dependent men: Impact of childhood emotional abuse and dissociation, Psychiatry Res. 186 (2011) 85–90. doi:10.1016/j.psychres.2010.07.004.

[28] K.M. Keyes, M.L. Hatzenbuehler, D.S. Hasin, Stressful life experiences, alcohol consumption, and alcohol use disorders: The epidemiologic evidence for four main types of stressors, Psychopharmacology (Berl). 218 (2011) 1–17. doi:10.1007/s00213-011-2236-1.

[29] L. Khoury, Y.L. Tang, B. Bradley, J.F. Cubells, K.J. Ressler, Substance use, childhood traumatic experience, and Posttraumatic Stress Disorder in an urban civilian population, Depress. Anxiety. 27 (2010) 1077–1086. doi:10.1002/da.20751.

[30] M.F. Lopez, R.I. Anderson, H.C. Becker, Effect of different stressors on voluntary ethanol intake in ethanol-dependent and nondependent C57BL/6J mice, Alcohol. 51 (2016) 17–23. doi:10.1016/j.alcohol.2015.11.010.

[31] R.H. Pietrzak, R.B. Goldstein, S.M. Southwick, B.F. Grant, Psychiatric comorbidity of full and partial posttraumatic stress disorder among older adults in the United States: Results from wave 2 of the national epidemiologic survey on alcohol and related conditions, Am. J. Geriatr. Psychiatry. 20 (2012) 380–390. doi:10.1097/JGP.0b013e31820d92e7.

[32] B.F. Grant, R.B. Goldstein, T.D. Saha, S. Patricia Chou, J. Jung, H. Zhang, R.P. Pickering, W. June Ruan, S.M. Smith, B. Huang, D.S. Hasin, Epidemiology of DSM-5 alcohol use disorder results from the national epidemiologic survey on alcohol and related conditions III, JAMA Psychiatry. 72 (2015) 757–766. doi:10.1001/jamapsychiatry.2015.0584.

[33] E.F.G. Naninck, P.J. Lucassen, J. Bakker, Sex Differences in Adolescent Depression: Do Sex Hormones Determine Vulnerability?, J. Neuroendocrinol. 23 (2011) 383–392. doi:10.1111/j.1365-2826.2011.02125.x.

[34] S. Nolen-Hoeksema, J.S. Girgus, The emergence of gender differences in depression during adolescence., Psychol. Bull. 115 (1994) 424–443. doi:10.1037/0033-2909.115.3.424.

[35] M. Piccinelli, G. Wilkinson, Gender differences in depression. Critical review, Br. J. Psychiatry. 177 (2000) 486–492. doi:10.1192/bjp.177.6.486.

[36] M. Steiner, E. Dunn, L. Born, Hormones and mood: from menarche to menopause and beyond., J. Affect. Disord. 74 (2003) 67–83. http://www.ncbi.nlm.nih.gov/pubmed/12646300.

[37] T.J. Wade, J. Cairney, D.J. Pevalin, Emergence of Gender Differences in Depression during Adolescence: National Panel Results from Three Countries, J. Am. Acad. Child Adolesc. Psychiatry. 41 (2002) 190–198. doi:10.1097/00004583-200202000-00013.

[38] J. Burgado, C.S. Harrell, D. Eacret, R. Reddy, C.J. Barnum, M.G. Tansey, A.H. Miller, H. Wang, G.N. Neigh, Two weeks of predatory stress induces anxiety-like behavior with co-morbid depressive-like behavior in adult male mice, Behav. Brain Res. 275 (2014) 120–125. doi:10.1016/j.bbr.2014.08.060.

[39] L.N. Eidson, M.E. deSousa Rodrigues, M.A. Johnson, C.J. Barnum, B.J. Duke, Y. Yang, J. Chang, S.D. Kelly, M. Wildner, R.J. Tesi, M.G. Tansey, Chronic psychological stress during adolescence induces sex-dependent adulthood inflammation, increased adiposity, and abnormal behaviors that are ameliorated by selective inhibition of soluble tumor necrosis factor with XPro1595, Brain. Behav. Immun. (2019). doi:10.1016/j.bbi.2019.06.027.

[40] C.J. Barnum, T.W.W. Pace, F. Hu, G.N. Neigh, M.G. Tansey, Psychological stress in adolescent and adult mice increases neuroinflammation and attenuates the response to LPS challenge, J. Neuroinflammation. 9 (2012). doi:10.1186/1742-2094-9-9.

[41] M.D. Casement, D.S. Shaw, S.L. Sitnick, S.C. Musselman, E.E. Forbes, Life stress in adolescence predicts early adult reward-related brain function and alcohol dependence, Soc. Cogn. Affect. Neurosci. 10 (2013) 416–423. doi:10.1093/scan/nsu061.

[42] C.E. Page, L. Coutellier, Adolescent Stress Disrupts the Maturation of Anxiety-related Behaviors and Alters the Developmental Trajectory of the Prefrontal Cortex in a Sex- and Age-specific Manner, Neuroscience. 390 (2018) 265–277. doi:10.1016/j.neuroscience.2018.08.030.

[43] S.A. Rowson, M. Bekhbat, S.D. Kelly, E.B. Binder, M.M. Hyer, G. Shaw, M.A. Bent, G. Hodes, G. Tharp, D. Weinshenker, Z. Qin, G.N. Neigh, Chronic adolescent stress sex-specifically alters the hippocampal transcriptome in adulthood, Neuropsychopharmacology. 44 (2019) 1207–1215. doi:10.1038/s41386-019-0321-z.

[44] M. Bekhbat, P.A. Howell, S.A. Rowson, S.D. Kelly, M.G. Tansey, G.N. Neigh, Chronic adolescent stress sex-specifically alters central and peripheral neuro-immune reactivity in rats, Brain. Behav. Immun. 76 (2019) 248–257. doi:10.1016/j.bbi.2018.12.005.

[45] S.A. Rowson, C.S. Harrell, M. Bekhbat, A. Gangavelli, M.J. Wu, S.D. Kelly, R. Reddy, G.N. Neigh, Neuroinflammation and behavior in HIV-1 transgenic rats exposed to chronic adolescent stress, Front. Psychiatry. 7 (2016) 102. doi:10.3389/fpsyt.2016.00102.

[46] L.N. Eidson, M.E. deSousa Rodrigues, M.A. Johnson, C.J. Barnum, B.J. Duke, Y. Yang, J. Chang, S.D. Kelly, M. Wildner, R.J. Tesi, M.G. Tansey, Chronic psychological stress during adolescence induces sex-dependent adulthood inflammation, increased adiposity, and abnormal behaviors that are ameliorated by selective inhibition of soluble tumor necrosis factor with XPro1595, Brain. Behav. Immun. (2019). doi:10.1016/j.bbi.2019.06.027.

[47] K.R. Urban, E. Geng, S. Bhatnagar, R.J. Valentino, Age- and sex-dependent impact of repeated social stress on morphology of rat prefrontal cortex pyramidal neurons, Neurobiol. Stress. 10 (2019) 100165. doi:10.1016/j.ynstr.2019.100165.

[48] S. Ordaz, B. Luna, Sex differences in physiological reactivity to acute psychosocial stress in adolescence, Psychoneuroendocrinology. 37 (2012) 1135–1157. doi:10.1016/j.psyneuen.2012.01.002.

[49] E.I. Varlinskaya, E.U. Kim, L.P. Spear, Chronic intermittent ethanol exposure during adolescence: Effects on stress-induced social alterations and social drinking in adulthood, Brain Res. 1654 (2017) 145–156. doi:10.1016/j.brainres.2016.03.050.

[50] L.E. Chaby, M.J. Sheriff, A.M. Hirrlinger, V.A. Braithwaite, Does early stress prepare individuals for a stressful future? Stress during adolescence improves foraging under threat, Anim. Behav. 105 (2015) 37–45. doi:10.1016/j.anbehav.2015.03.028.

[51] M. Bardi, C.L. Franssen, J.E. Hampton, E.A. Shea, A.P. Fanean, K.G. Lambert, Paternal experience and stress responses in California mice (Peromyscus californicus)., Comp. Med. 61 (2011) 20–30. http://www.ncbi.nlm.nih.gov/pubmed/21819678%0Ahttp://www.pubmedcentral.nih.gov/articlerender.fcgi?artid=PMC3060428

[52] S.L. Byers, M. V. Wiles, S.L. Dunn, R.A. Taft, Mouse estrous cycle identification tool and images, PLoS One. 7 (2012) e35538. doi:10.1371/journal.pone.0035538.

[53] V. Carola, F. D’Olimpio, E. Brunamonti, F. Mangia, P. Renzi, Evaluation of the elevated plus-maze and open-field tests for the assessment of anxiety-related behaviour in inbred mice, Behav. Brain Res. 134 (2002) 49–57. doi:10.1016/S0166-4328(01)00452-1.

[54] E. Choleris, A.W. Thomas, M. Kavaliers, F.S. Prato, A detailed ethological analysis of the mouse open field test: Effects of diazepam, chlordiazepoxide and an extremely low frequency pulsed magnetic field, Neurosci. Biobehav. Rev. 25 (2001) 235–260. doi:10.1016/S0149-7634(01)00011-2.

[55] L. Prut, C. Belzung, The open field as a paradigm to measure the effects of drugs on anxiety-like behaviors: A review, Eur. J. Pharmacol. 463 (2003) 3–33. doi:10.1016/S0014-2999(03)01272-X.

[56] L.S. Hwa, A. Chu, S.A. Levinson, T.M. Kayyali, J.F. Debold, K.A. Miczek, Persistent escalation of alcohol drinking in C57BL/6J mice with intermittent access to 20% ethanol, Alcohol. Clin. Exp. Res. 35 (2011) 1938–1947. doi:10.1111/j.1530-0277.2011.01545.x.

[57] F.W. Hopf, S.J. Chang, D.R. Sparta, M.S. Bowers, A. Bonci, Motivation for alcohol becomes resistant to quinine adulteration after 3 to 4 months of intermittent alcohol self-administration, Alcohol. Clin. Exp. Res. 34 (2010) 1565–1573. doi:10.1111/j.1530-0277.2010.01241.x.

[58] D.K. Cozzoli, M.A. Tanchuck-Nipper, M.N. Kaufman, C.B. Horowitz, D.A. Finn, Environmental stressors influence limited-access ethanol consumption by C57BL/6J mice in a sex-dependent manner, Alcohol. 48 (2014) 741–754. doi:10.1016/j.alcohol.2014.07.015.

[59] L.D. Middaugh, B.M. Kelley, Operant Ethanol Reward in C57BL/6 Mice, Alcohol. 17 (1999) 185–194. doi:10.1016/s0741-8329(98)00056-1.

[60] L.D. Middaugh, B.M. Kelley, A.-L.E. Bandy, K.K. McGroarty, Ethanol Consumption by C57BL/6 Mice, Alcohol. 17 (1999) 175–183. doi:10.1016/s0741-8329(98)00055-x.

[61] F. Faul, E. Erdfelder, A.-G. Lang, A. Buchner, G*Power 3: a flexible statistical power analysis program for the social, behavioral, and biomedical sciences., Behav. Res. Methods. 39 (2007) 175–91. http://www.ncbi.nlm.nih.gov/pubmed/17695343.

[62] K. Goltseker, F.W. Hopf, S. Barak, Advances in behavioral animal models of alcohol use disorder, Alcohol. (2018). doi:10.1016/j.alcohol.2018.05.014.

[63] D. Darevsky, T.M. Gill, K.R. Vitale, B. Hu, S.A. Wegner, F.W. Hopf, Drinking despite adversity: behavioral evidence for a head down and push strategy of conflict-resistant alcohol drinking in rats, Addict. Biol. 24 (2019) 426–437. doi:10.1111/adb.12608.

[64] E.A. Sneddon, R.D. White, A.K. Radke, Sex Differences in {Binge-Like} and {Aversion-Resistant} Alcohol Drinking in {C57BL/6J} Mice, (2018). doi:10.1111/acer.13923.

[65] J.J. Olney, S.A. Marshall, T.E. Thiele, Assessment of depression-like behavior and anhedonia after repeated cycles of binge-like ethanol drinking in male C57BL/6J mice, Pharmacol. Biochem. Behav. 168 (2018) 1–7. doi:10.1016/j.pbb.2018.03.006.

[66] R. Adamec, A. Holmes, J. Blundell, Vulnerability to lasting anxiogenic effects of brief exposure to predator stimuli: Sex, serotonin and other factors-Relevance to PTSD, Neurosci. Biobehav. Rev. 32 (2008) 1287–1292. doi:10.1016/j.neubiorev.2008.05.005.

[67] M.J. Caruso, L.R. Seemiller, T.B. Fetherston, C.N. Miller, D.E. Reiss, S.A. Cavigelli, H.M. Kamens, Adolescent social stress increases anxiety-like behavior and ethanol consumption in adult male and female C57BL/6J mice, Sci. Rep. 8 (2018) 10040. doi:10.1038/s41598-018-28381-2.

[68] K.M. Lee, M. Coehlo, H.A. McGregor, R.S. Waltermire, K.K. Szumlinski, Binge alcohol drinking elicits persistent negative affect in mice, Behav. Brain Res. 291 (2015) 385–398. doi:10.1016/j.bbr.2015.05.055.

